# Ion Mobility Mass Spectrometry Guided Modeling with AlphaFold and Rosetta Improves Protein Complex Structure Prediction

**DOI:** 10.64898/2026.02.16.706193

**Authors:** Akshaya Narayanasamy, Zachary C. Drake, S. M. Bargeen A. Turzo, Amber D. Rolland, James S. Prell, Vicki H. Wysocki, Steffen Lindert

**Affiliations:** Department of Chemistry and Biochemistry, University of California, Los Angeles, CA, 90095, USA; Center for Computational Biology, Flatiron Institute, 162 Fifth Avenue, New York, NY, 10010, USA; Department of Chemistry and Biochemistry and Materials Science Institute, 1253 University of Oregon, Eugene, OR, 97403, USA; School of Chemistry and Biochemistry, Georgia Tech, Atlanta, GA, 30332, USA

## Abstract

Ion mobility mass spectrometry (IM-MS) provides valuable structural information about protein shape and size through collision cross section (CCS). However, it lacks atomic level structural detail. While AlphaFold has been successful in predicting monomeric protein structure, it can struggle with modeling protein complexes. To address these limitations, we developed a method that integrates IM-MS data with AlphaFold and Rosetta to improve complex structure prediction. Our approach uses experimental CCS data to guide the assembly of AlphaFold predicted subunits using a Rosetta docking pipeline and evaluating the resulting complexes with a newly developed score. Using this strategy, we were able to improve root mean square deviation (RMSD) values for 26 of 38 (68%) complexes compared to AlphaFold-Multimer. Furthermore, 16 of these systems improved significantly from greater than 4 Å RMSD to less than 4 Å. This method demonstrates a robust approach to overcome limitations in complex assembly modeling.

**Table of Contents Graphic:** In this integrative modeling work, protein complex structures were modeled by combining AlphaFold predicted subunits with Rosetta docking. Collision cross section data from ion-mobility mass spectrometry were used as evaluation constraints and docked models were scored using the IM-complex score. The best scoring models generally represent accurate protein complex structures.

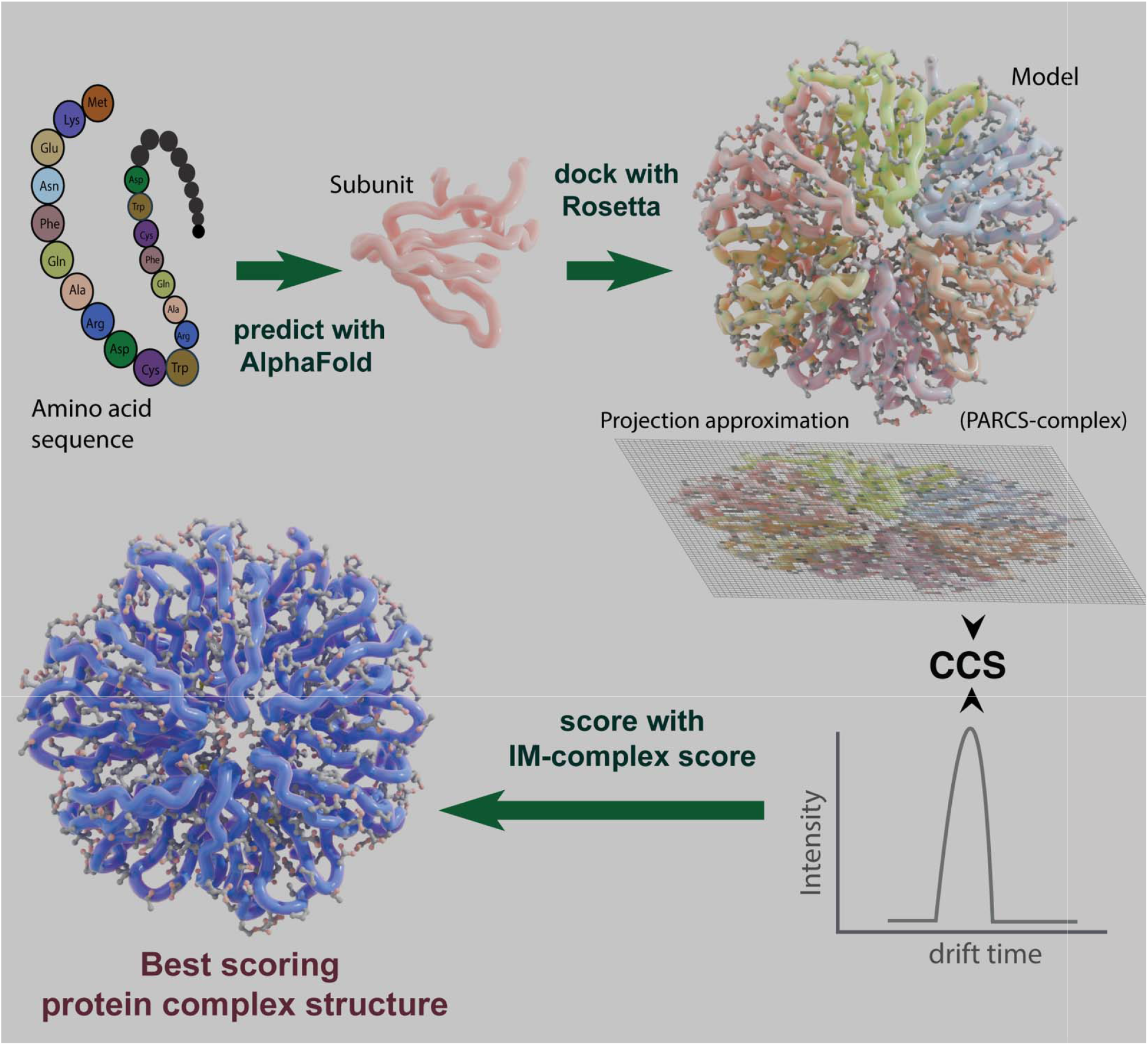

## Introduction

The intricate nature of cellular functions depends on the interaction and coordination of multiple protein subunits that form protein complexes.^1,2^ As these protein complexes are essential for many important biological processes, elucidating the structure of protein complexes is crucial for understanding disease mechanisms and advancing the development of therapeutic treatments.^3^ Only a small fraction of the human proteome has been structurally resolved experimentally and structural information on protein complexes is even more limited.^4^ Traditional experimental approaches for obtaining protein structures such as X-ray crystallography, nuclear magnetic resonance spectroscopy, and cryo-electron microscopy each present challenges related to protein size, crystallization requirements, and achievable resolution. Additionally, these methods are time consuming and resource intensive.^5,6^ This represents a major bottleneck in biomedical research and drug discovery, highlighting the need for alternative approaches to extract structural information.

Native mass spectrometry has emerged as a complementary approach for analyzing proteins and their complexes in a folded, native state, providing insights into protein-protein interactions.^7^ These experiments have high sensitivity, low sample requirement, no size constraint, can be performed on mixtures and are less time-intensive, making them suitable for high-throughput analysis.^7–10^ In addition to these advantages, native mass spectrometry encompasses several techniques that together provide a broad range of structural information. For example, native top-down mass spectrometry provides detailed analysis of protein composition,^11^ ion mobility reveals protein size and shape^12^, collision-induced dissociation examines complex stability, and surface-induced dissociation identifies subunit connectivity.^13^

In particular, Ion Mobility Mass Spectrometry (IM-MS) is a powerful analytical technique for characterizing and separating ions based on size and shape (as interpreted from their rotationally-averaged collisional cross-sections). In IM-MS, protein ions are transmitted through to a neutral buffer gas in a drift tube.^14^ They undergo consecutive collisions with the neutral gas atoms/molecules, leading to different drift velocities based on their size, shape and charge before reaching the mass analyzer and detector. The average momentum exchanged between an ion and the surrounding gas particles is often estimated using the momentum transfer integral and can be interpreted in terms of the ion’s collision cross section (CCS). CCS measures the effective cross-sectional area of a protein ion and gas molecules during collisions, reflecting the overall shape and size of a protein complex. Since different protein conformations present different effective areas, CCS measurements can be used to distinguish complexes based on their three-dimensional shape.^12,15–17^

Computational techniques provide an alternative pathway for obtaining protein structures in cases where traditional experimental methods are not viable. DeepMind’s AlphaFold^18–20^ (AF) has revolutionized computational protein structure modeling through deep learning, driving transformative advances in structural biology and contributing to the scientific progress recognized by the 2024 Nobel Prize in Chemistry. Alongside AF, methods such as RoseTTAFold^21,22^ and Boltz^23,24^ have also significantly impacted medical and biotechnological research.^25^ Despite their successes, deep learning methods face limitations when predicting the structures of certain types of proteins, such as protein complexes or membrane-bound proteins, proteins with few known homologous sequences, intrinsically disordered proteins, proteins undergoing dynamic motions,^26^ and ternary complexes.^27^ Methods like AF have advanced protein structure prediction by leveraging coevolutionary signals from multiple sequence alignments (MSAs); however, applying these approaches to protein complexes proves to be significantly more challenging. In homo-oligomers, identical MSAs make it difficult to distinguish signals between chains, while in hetero-oligomers, sequence pairing weakens the signal, complicating detection.^28^ Previous benchmark studies show that AlphaFold-Multimer (AFM) exhibits an accuracy of ~ 43% – 70 % in predicting protein complexes^19,29^, indicating a need for improvement. In this work, we focused on developing a method to improve structure prediction specifically for protein complexes with the aid of IM-MS data.

Conventional protein-protein docking methods are valuable for predicting and analyzing complex structures. In these approaches, individual monomers obtained through computational or experimental means are assembled into complexes by sampling possible conformations and identifying the most energetically favorable arrangement. State-of-the-art protein-protein docking tools include EvoDock,^30^ AlphaRED,^31^ ClusPro,^32^ HDOCK,^33^ ZDOCK,^34^ Rosetta Symmetric Docking^35–37^ and RosettaDock.^38–40^ The Rosetta software suite offers a wide range of computational algorithms for the modeling and analysis of protein structures. RosettaDock and Symmetric Docking are part of the Rosetta software suite which utilizes Monte Carlo simulations to generate docked protein complexes from subunits.

Previous studies have shown that integrating mass spectrometry data with the Rosetta modeling platform can improve protein structure prediction.^41–60^ In prior work we have incorporated IM-MS data into Rosetta and have shown that this approach is successful in accurately predicting monomeric protein structures.^50^ In this work, we adopted an integrative modeling approach by incorporating sparse CCS data from IM-MS experiments with AF and Rosetta to enhance the prediction of protein complex structures. We developed an updated version of PARCS (PARCS-complex) that can simulate CCS values for protein complexes with mean absolute percent error of only 3.9 %. We further developed an additional score, ‘IM-complex score’, that integrated IM-MS data into the Rosetta interface score (Rosetta Isc). We validated the score function on datasets with simulated CCS values and with experimental CCS measurements, respectively. Use of the IM-complex score improved ranking of proteins in both datasets. When tested on an exclusive dataset with poor AFM predictions, our method exhibited an improvement for 26 out of 38 protein complexes. Among those, 19 of the modeled protein complexes had RMSD□□□ (root mean square deviation normalized over sequence length) less than 4 Å, indicating native-like structures. The PARCS-complex and IM-complex score function applications are available as a part of the Rosetta suite, and a detailed description of their usage is given in the supplementary information.

## Results and Discussion

### Original PARCS underestimated actual CCS values for protein complexes

In previous work,^50^ we introduced the Projection Approximation using Rough Circular Shapes (PARCS), an adaptation of the projection approximation (PA) method implemented in the Rosetta molecular modeling suite. PARCS works by projecting the 3D protein structure onto a 2D grid and estimating the projection area using a 9-point circular approximation for each atom. This process is repeated across multiple random orientations, and the average projection area is used to generate the final CCS prediction.^61^

While PARCS demonstrated high accuracy in predicting CCS values for monomeric proteins, tests on protein complexes with available experimental CCS values showed that the prediction error increased for these larger assemblies. PARCS consistently underestimated experimental CCS values, as shown in Figure 1a, with mean absolute percent error (MAPE) of 8.1%. This systematic underestimation was expected since the PA method does not account for scattering and long-range interactions during collisions, leading to oversimplified results, with particularly notable deviations for larger proteins.^62^

**Figure 1.**
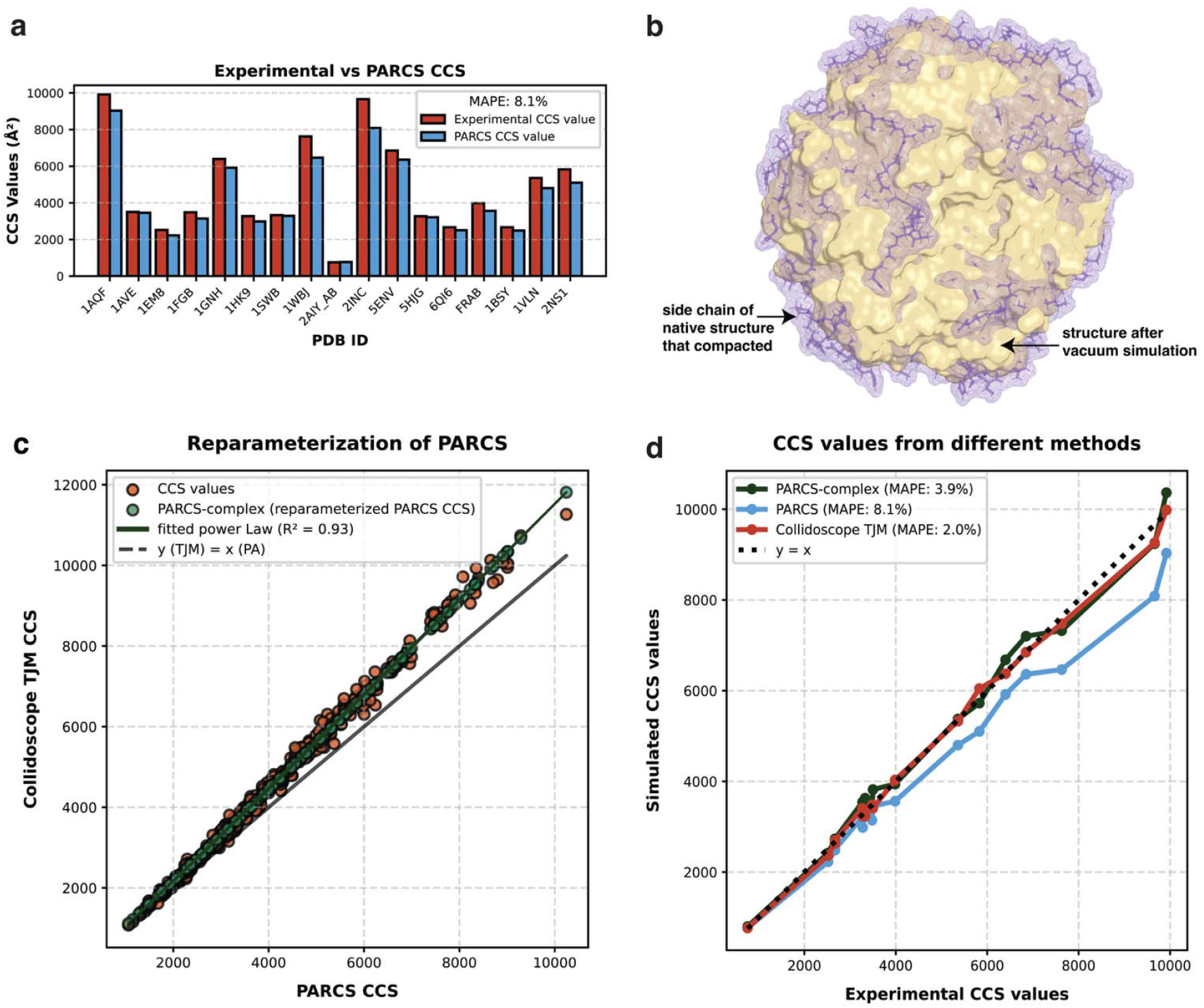
Reparameterization of PARCS. a) Comparison of experimental CCS values and PARCS CCS values for 17 protein complexes. The red bars represent the experimental CCS values and blue bars represent predicted PARCS CCS values. Mean absolute percent error (MAPE) was used to calculate errors. b) Overlay of protein structure before and after vacuum MD simulation. Purple mesh with highlighted sidechains is the native protein structure (Hfq protein from Escherichia coli, PDB ID: 1HK9) while the yellow-colored surface is the same protein after side chain compaction following vacuum simulation. c) The orange points are the Collidoscope CCS values plotted against the PARCS CCS values. Green points are the PARCS-complex (reparametrized PARCS CCS) values. d) Comparison of CCS values simulated by different methods to experimental CCS values of 17 protein complexes. Blue points are PARCS CCS values, orange-colored points are Collidoscope TJM CCS values, green points are PARCS-complex CCS values.

Although the PA has limitations when applied to protein complexes, studies have shown that it remains helpful in estimating CCS values by correlating PA-derived results with those obtained from the Trajectory Method (TJM).^62,63^ Unlike the PA, the TJM approach accounts for factors such as long-range interactions, multiple scattering events, ion charge, and temperature effects, resulting in more accurate predicted CCS values.^17^ However, this increased TJM accuracy comes at the cost of significantly greater computational time, being approximately 10□ times slower than the PA.^63^ By reparametrizing the PA to yield results closer to TJM CCS values, it is possible to leverage the computational efficiency of PA while taking advantage of more accurate TJM predictions.^62,63^

### TJM CCS calculations for proteins in the reparameterization dataset

To investigate the relationship between PARCS and TJM CCS values, we analyzed a dataset of 461 protein complexes (see Methods, Reparameterization dataset). TJM CCS values were calculated using the Collidoscope^64^ software, following its established protocol.^65^ In this approach, the gas-phase environment of IM-MS was simulated via vacuum simulations to generate protein structures suitable for TJM CCS estimation. Because this low-pressure environment lacks solvent molecules, charged groups undergo self-solvation, resulting in the collapse of the side chains of protein complexes.^65^ Structures of one protein complex (Hfq protein from Escherichia coli, PDB ID: 1HK9) from the benchmark dataset before and after MD simulations are shown in Figure 1b. This clearly depicts the side chains (highlighted in purple color) collapsing while simultaneously maintaining the protein tertiary backbone structure (yellow colored surface). The average normalized radius of gyration changed by 0.070 Å across 461 complexes, indicating minor backbone structural changes (see Figure S1). With these compact structures, TJM CCS values were computed with Collidoscope to achieve an approximate experimental accuracy. For comparison, PARCS CCS values were calculated in parallel within the Rosetta suite directly from the experimental PDB structures of all proteins in the reparameterization dataset.

### Reparameterization of PARCS with TJM CCS values improved CCS values for protein complexes

We found that TJM Collidoscope CCS values and PARCS CCS values were strongly correlated, with an R^2^ of 0.93. As seen from the deviation of the orange points from the dashed line in Figure 1c, TJM yielded significantly higher CCS values than PA, particularly for larger proteins. The relationship between TJM Collidoscope CCS values and PARCS CCS values could be described by a power law (Eqn. 1) The reparameterized PARCS is referred to as PARCS-complex.

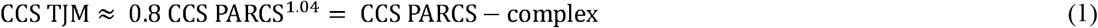

This reparameterization allowed PARCS-complex results to have greater alignment with TJM values, resulting in a MAPE of only 2.5 % (as opposed to 6.3% without) between the two methods (see alignment of green and orange points in Figure 1c). We validated our approach on a benchmark dataset of proteins with known experimental CCS values as shown in Figure1d and observed that the PARCS-complex error decreased from 8.1% to 3.9% when using the reparametrized version of PARCS (see Figure 1d). The PARCS-complex algorithm was comparably accurate to Collidoscope TJM (MAPE of 2.0%) in predicting CCS data, while retaining the speed of PA methods for protein complexes. Overall, this reparameterization improved the accuracy of PARCS, leading to more reliable predictions of CCS for protein complexes, and thus enabling the use of PARCS-complex in structure prediction scenarios.

### IM-complex score outperformed Rosetta Isc to identify native-like structures with ideal CCS data

Next, we developed an IM-complex score function that uses CCS values from PARCS-complex to improve scoring of predicted assemblies. In previous work, we developed an IM score function for monomer structure prediction, which performed well in identifying native-like monomeric structures^50^. However, that function was not applicable to complexes, as protein complexes are larger in size and have greater variation in CCS values compared to monomeric structures. To address this, we developed a protein complex specific IM-complex score function in Rosetta (see Methods, Eqn. 2) to evaluate the agreement between predicted models and CCS data from IM-MS.

To validate the IM-complex score function, we assembled an ideal dataset with 38 challenging protein complexes (ideal dataset) for which AFM failed to predict accurate quaternary structures (RMSD > 4 Å). We analyzed AFM confidence metrics of these tough targets, including pLDDT (predicted Local Distance Difference Test) and the predicted model confidence score, and compared them against RMSD values calculated with respect to the corresponding ground-truth native PDB structures. Notably, despite poor structural accuracy (every RMSD > 4 Å), 22 complexes exhibited high average pLDDT scores (≥ 90), and 13 complexes showed high predicted model confidence (≥ 0.5) (see Figure S2). These results demonstrate that AFM confidence metrics provide potentially inaccurate estimates of the accuracies of predicted structural coordinates.^26,66,67^

This ‘ideal dataset’ included both homo- and hetero-oligomeric assemblies, spanning stoichiometries from dimers to hexamers (2–6 subunits) and total sequence lengths ranging from 102 to 2305 amino acids (see Methods, ideal dataset). Since no IM-MS data was available for this dataset, we used PARCS-complex to simulate CCS values based on the experimental reference structures. The monomeric chains of the predicted subunits showed an average RMSD of 1.04 Å, ensuring that the subunit conformations were accurate representations (see Table S1). After docking the subunits with Rosetta, we next scored the generated models with the IM-complex score and Rosetta Isc function to assess whether IM-MS data can provide additional shape information to aid native-like structure identification. As shown in Figure 2a and b, the IM-complex score function improved recognition of near-native structures. The average RMSD of the ideal dataset proteins decreased from 7.0 Å without the IM-complex score to 6.0 Å with the IM-complex score. The improvement was also quantified using the Template Modelling score (TM-score), a standard metric that measures how similar the fold of a predicted structure is to a reference structure, where values closer to 1 indicate better agreement. The increase in the average TM-score from 0.64 to 0.69 therefore reflected more accurate global folds of the proteins. Of the 38 systems, Rosetta Isc identified only 14 systems with RMSD□□□ less than 4.0 Å (corresponding to near-native complexes), whereas IM-complex score enabled accurate modeling of 19 systems. A similar trend was observed in TM-score analyses (see Figure 2b). Of the 38 protein complexes, 11 showed improvement in quality metrics after rescoring with the IM-complex score (Eq. 3 in Methods) with an average reduction in RMSD□□□ of 4.6 Å and an average increase in TM-score of 0.15 for the proteins that improved in structure quality.

**Figure 2.**
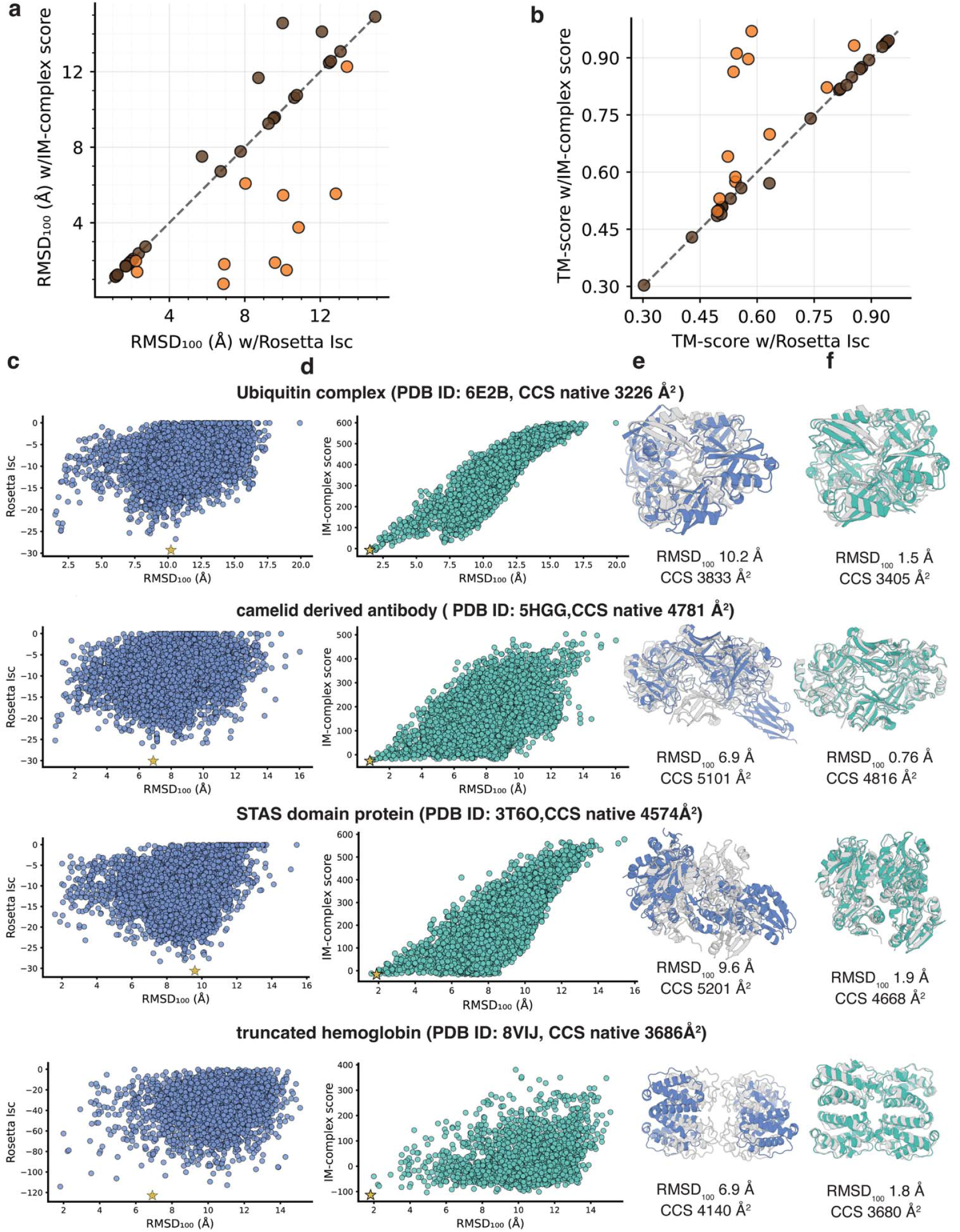
Improved model selection was observed when using the IM-complex score for the 38 proteins in the ideal dataset. The predicted models from the IM-complex score were compared to those scored with the Rosetta Isc function in terms of their a) RMSD□□□ (root mean square deviation normalized by sequence length) and b) TM-Score (template modeling score). The protein complex structures that improved upon using the IM-complex score are colored orange, while the remaining proteins are colored brown. c-f) Comparison of predicted structures of four significantly improved proteins with CCS values calculated from their PDB structure using PARCS-complex. Respective protein score vs RMSD□□□ plots scored with c) Rosetta Isc (blue scatter plot) and d) IM-complex score (green scatter plot). The best scoring model is marked with a yellow star in each plot. Corresponding best scoring structures generated with e) the Rosetta Isc function (blue) and f) the IM-complex score function (green) are compared to their native structures (gray) and respective RMSD□□□ and PARCS-complex CCS values are listed.

Four proteins (ubiquitin complex (PDB ID: 6E2B, homo 6-mer), camelid derived antibody (PDB ID: 5HGG, hetero 4-mer), structure of an anti-sigma-factor antagonist (STAS) domain protein (PDB ID: 3T6O, homo 6-mer), and truncated hemoglobin (PDB ID: 8VIJ, homo 4-mer)) showed significant improvement in RMSD□□□ (from greater than 6 Å to less than 3 Å) after rescoring with the IM-complex score. Analysis of these four complexes (see Figure 2c) showed that Rosetta was capable of generating near-native conformations; however, the Rosetta Isc often failed to rank them correctly. Scoring with IM-complex score introduced complementary shape constraints, enabling more accurate selection of native-like poses (Figure 2d). We compared the native structure to the best scored model using Rosetta Isc (Figure 2e) and the IM-complex score function (Figure 2f), respectively. Notably, the best-scoring models showed improvement in accuracy and also showed simulated collision cross sections (CCS) that more closely matched the native values (see labels of Figure 2e and 2f), highlighting the effectiveness of the IM-complex score function.

### IM-complex integrative modeling method accurately modeled many protein complexes that are poorly predicted by AFM

Next, we compared the best scoring structures identified by the IM-complex score function with those predicted by AFM. Generally, structures modeled with the IM-complex pipeline had better RMSD□□□ and TM-scores than models predicted by AFM in the ideal dataset as shown in Figure 3a. Specifically, RMSD□□□ values for 26 of the 38 protein complexes improved, with an average decrease in RMSD□□□ values of 6.4 Å. A similar trend was observed for the TM-score (Figure 3b), where 23 of the 38 complexes showed better scores, with an average TM-score increase of 0.3 among the improved proteins.

**Figure 3.**
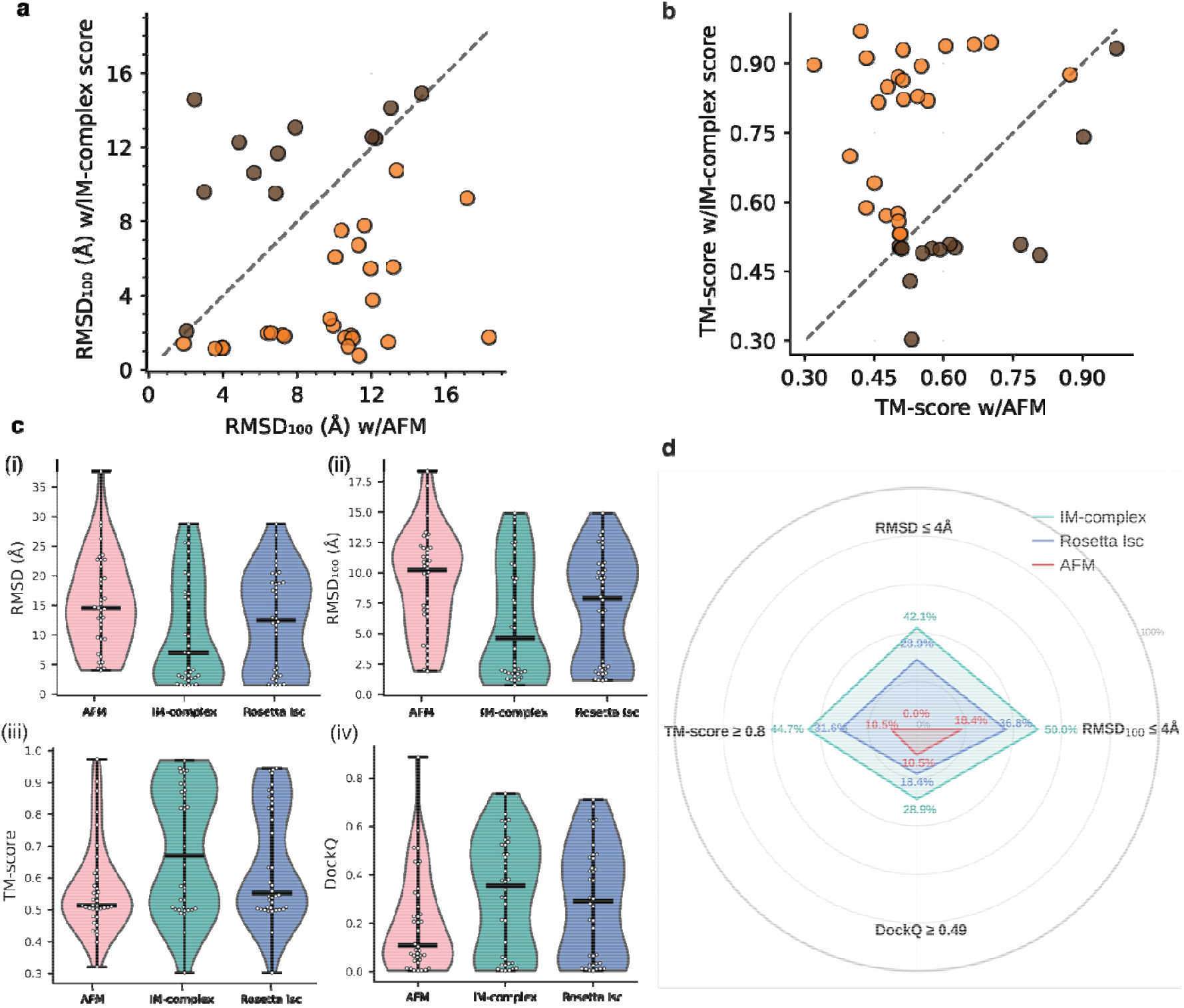
Comparison of structures predicted for the ideal dataset to those generated with AFM. Comparison of structures modeled using the IM-complex docking pipeline and AFM, using a) RMSD□□□ (root mean square deviation normalized by sequence length) and b) TM-score (template modeling score). The protein omplex structures that improved with the IM-complex score compared to AFM are colored orange, and all others are colored brown. c) Violin plots of structures modeled by three different modeling methods: AFM (pink), IM-complex score (green), Rosetta Isc (blue). The dark lines represent the median of the ideal dataset with each method. i) RMSD ii) RMSD□□□ iii) TM-Score, and iv) DockQ score. d) Radar plot comparing the performance of AFM, IM-complex score and Rosetta Isc predicted models across four evaluation metrics (RMSD, RMSD□□□, TM-score, DockQ score). Each axis on the plot represents stringent accuracy criteria, including RMSD ≤ 4 Å, RMSD□□□ ≤ 4 Å, TM-score ≥ 0.5, and DockQ ≥ 0.49, and the area enclosed by each polygon reflects the overall performance profile. Models with larger, more outward-reaching shapes perform better across multiple criteria. The green polygon is IM-complex score, the blue polygon is Rosetta Isc, and the pink is that of AFM models.

This trend of improvement among all 38 protein complexes, across different modeling methods (AFM, IM-complex score, and Rosetta Isc) was analyzed using different metrics and is highlighted in Figure 3c (i)-(iv). The median RMSD observed for AFM, Rosetta, and the IM-complex protocol was 14.5 Å, 12.5 Å and 7.0 Å respectively, indicating that structures modeled by the IM-complex protocol were more representative of native structures. Likewise, after normalizing the RMSD by the sequence length, the median RMSD□□□ of structures modeled by the IM-complex protocol (4.6 Å) was lower than that obtained with AFM (10.2 Å), and Rosetta (7.9Å) structures. In addition, median TM-scores were higher for complexes modeled with the IM-complex method (value of 0.67), compared to median values of 0.51 for AFM and 0.55 for Rosetta. This reflected a better global topology of complexes modeled with the IM-complex method. Finally, median DockQ scores were also highest for the IM-complex modeled complexes with a value of 0.36, compared to 0.10 for AFM and 0.30 for Rosetta, demonstrating clear gains in interface accuracy of complexes modeled with the IM-complex method.

The consolidated radar plot (Figure 3d) summarizes the comparative performance of AFM, Rosetta Isc, and the IM-complex score across the ideal dataset. The plot uses commonly accepted cutoffs for defining native-like structures: RMSD and RMSD□□□ values of 4 Å or less for native-like structures, TM-scores of at least 0.5 for good global folds, and DockQ values of 0.49 or higher for acceptable or high-quality models. The polygon for the IM-complex score spans the largest area, corresponding to an overall accuracy of roughly 28.9 – 50%. Rosetta Isc shows intermediate performance (18.4 – 36.8%), consistent with its ability to generate but not consistently rank near-native poses. In contrast, AFM covers the smallest area (0 – 18.4%), reflecting its lower accuracy on these challenging cases. Together, these results show that integrating experimental constraints from IM-MS data can help bridge the gap between physics-based docking and deep learning-based predictions, while outperforming AFM in challenging cases.

### IM-complex pipeline improved model quality for ideal dataset, frequently surpassing AFM

For several complexes in the ideal dataset, one of the main reasons for AFM failure was incorrect or distorted symmetry, which led to unrealistic overall shapes. One of the significantly improved protein complexes was the ubiquitin complex (PDB ID: 6E2B, homo 6-mer) (see Figure 4a). AFM produced an interface with a physically unreasonable large distance (33.1 Å) between chains that are experimentally observed to interact, disrupting the expected D3 symmetry and resulting in an incorrect overall assembly (RMSD of 22.7 Å). In contrast, assembling and scoring the subunits using the IM-complex score recovered the correct symmetry and produced a native-like structure (RMSD of 2.6 Å). A similar issue was observed for the dArc1 capsid (PDB ID: 6TAR, homo 5-mer) (see Figure 4b) and camelid derived antibody (PDB ID: 5HGG, hetero 4-mer) (see Figure 4C). AFM arranged the subunits in a linear configuration, losing the expected cyclic symmetry (C5 and C2, respectively), which led to inaccurate quaternary structures (RMSDs of 37.6 Å and 22.8 Å, respectively). Thus, although AF predicted the individual monomers with good accuracy (see Supplementary Table 1), it failed to reconstruct the correct multimeric assembly. Using Rosetta docking in combination with the IM-complex score restored the native-like symmetric arrangements for both complexes (RMSDs of 3.6 Å and 1.5 Å, respectively). For STAS domain protein (PDB ID: 3T6O, homo 6-mer) (see Figure 4d), the truncated hemoglobin complex (PDB ID: 8VIJ, homo 4-mer) (see Figure 4e), and A53 complex (PDB ID:8AVU, homo 2-mer) (see Figure 4f) AFM did not identify the correct interfaces, and their expected cyclic and dihedral symmetries (C2, D2, C2 respectively). These interactions were correctly modeled when using the IM-complex score, resulting in structures that matched the experimental quaternary arrangements.

**Figure 4.**
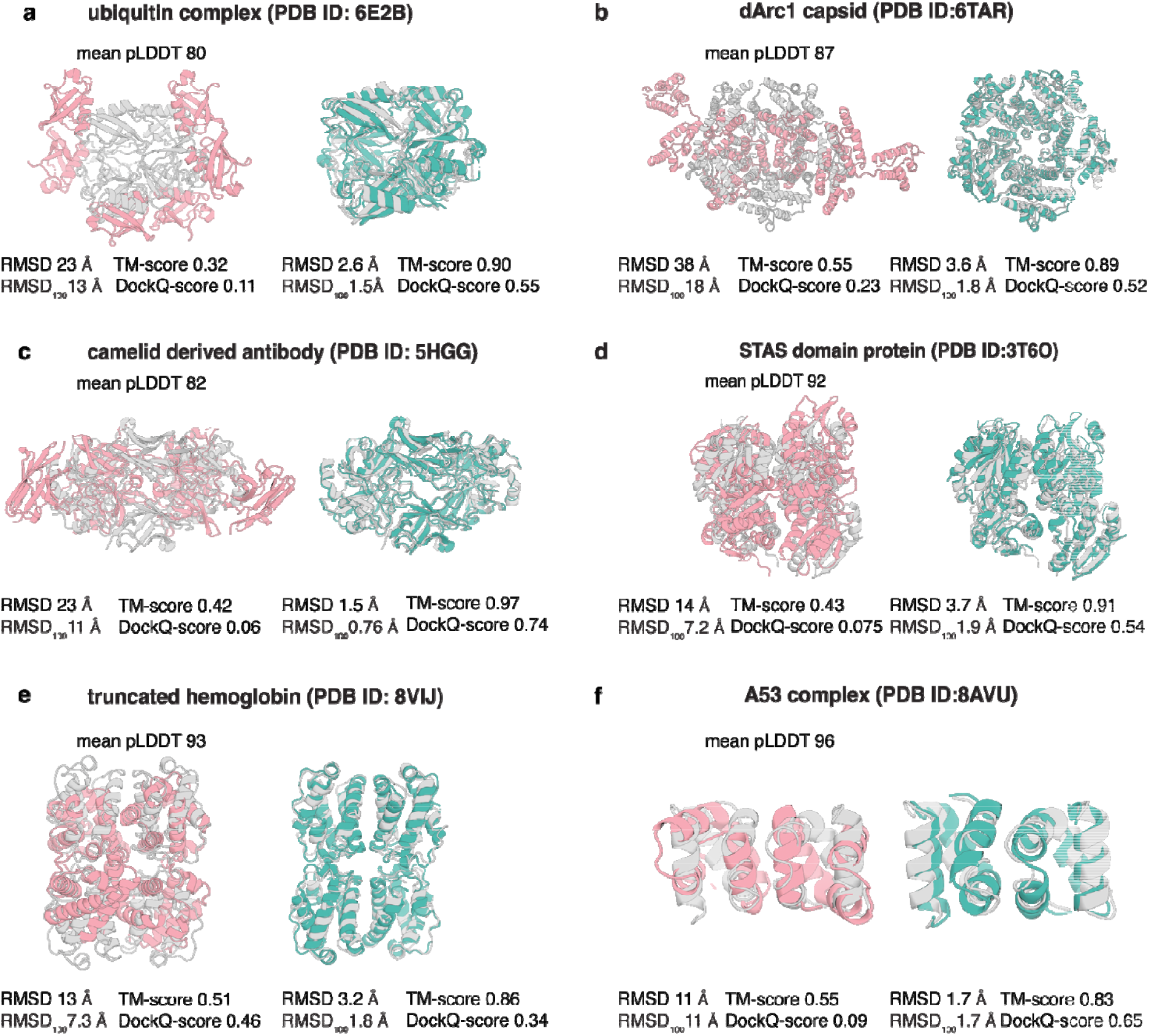
IM-complex pipeline improves modeling of protein complexes compared to AFM. For each protein complex (panels a–f), the AFM prediction is shown in pink, and the corresponding structure modeled using the IM-complex score is shown in green and native structures are shown in gray color.

This highlights a key advantage of incorporating IM-based shape information into structure selection. While AF can produce accurate monomeric subunits, it sometimes fails to infer correct symmetry in multimeric assemblies. Integrating physical shape constraints through the IM-complex score helps guide Rosetta to identify the proper arrangement, ensuring the correct overall shape and symmetry of the complex.

These improvements were not limited to a single type of complex, with 26 protein complexes showing improved RMSD across both homomers and heteromers. These improved complexes spanned all stoichiometries examined (2, 3, 4, 5, 6), with sequence lengths ranging from 102 to 2305 amino acids (refer to Table S1). This shows that the docking protocol with the IM-complex score function is applicable across different types of protein assemblies and is not restricted to a specific size or symmetry. Overall, the results clearly showed that adding sparse IM-MS data to the scoring process facilitated selecting native-like structures in situations where AFM alone struggled.

### IM-complex score function outperformed Rosetta Isc with experimental CCS data

We further evaluated the IM-complex score function using experimental CCS data. This was done to determine whether these measurements could improve structure selection compared to using the Rosetta Isc alone and to access how robust the score function is to experimental noise. Our benchmark dataset included 17 homomeric and heteromeric protein complexes. For all proteins in the benchmark dataset, the monomeric chains of the predicted subunits had an average RMSD of 1.25 Å (Table S2). We then generated 10,000 models per protein using the Rosetta docking pipeline. Finally, we scored all models with and without incorporating experimental CCS values to assess whether IM-MS data helped identify more native-like poses. Protein complexes in the benchmark dataset represent some of the most well-studied proteins in biochemistry. 15 of the 17 protein structures were deposited in the Protein Data Bank (PDB) before the year 2018 and were thus included in the AFM training set.^19^ Hence, a comparison to AFM is not meaningful in this case.

Notably, after rescoring, no protein in the dataset showed worse RMSD or RMSD□□□ values (Figure 5a), indicating that incorporating experimental IM-MS data is at least neutral and often beneficial for structure selection compared to using the Rosetta Isc alone. The IM-complex score provided meaningful improvements in ranking near-native poses generated by Rosetta. Before rescoring, 6 out of 17 complexes had RMSD□□□ values below 5 Å. After rescoring with the IM-complex score, 10 out of 17 complexes achieved RMSD□□□ values below 5 Å. The average RMSD□□□ and TM-score before rescoring was 7.32 Å and 0.61, respectively, whereas after rescoring they improved to 4.82 Å and 0.71. These shifts reflect a clear trend toward lower RMSD□□□ values and higher TM-scores. This improvement is visually apparent in the scatter plots, where points move toward the lower RMSD region (Figure 5a) and higher TM-score region (Figure 5b) after rescoring.

**Figure 5.**
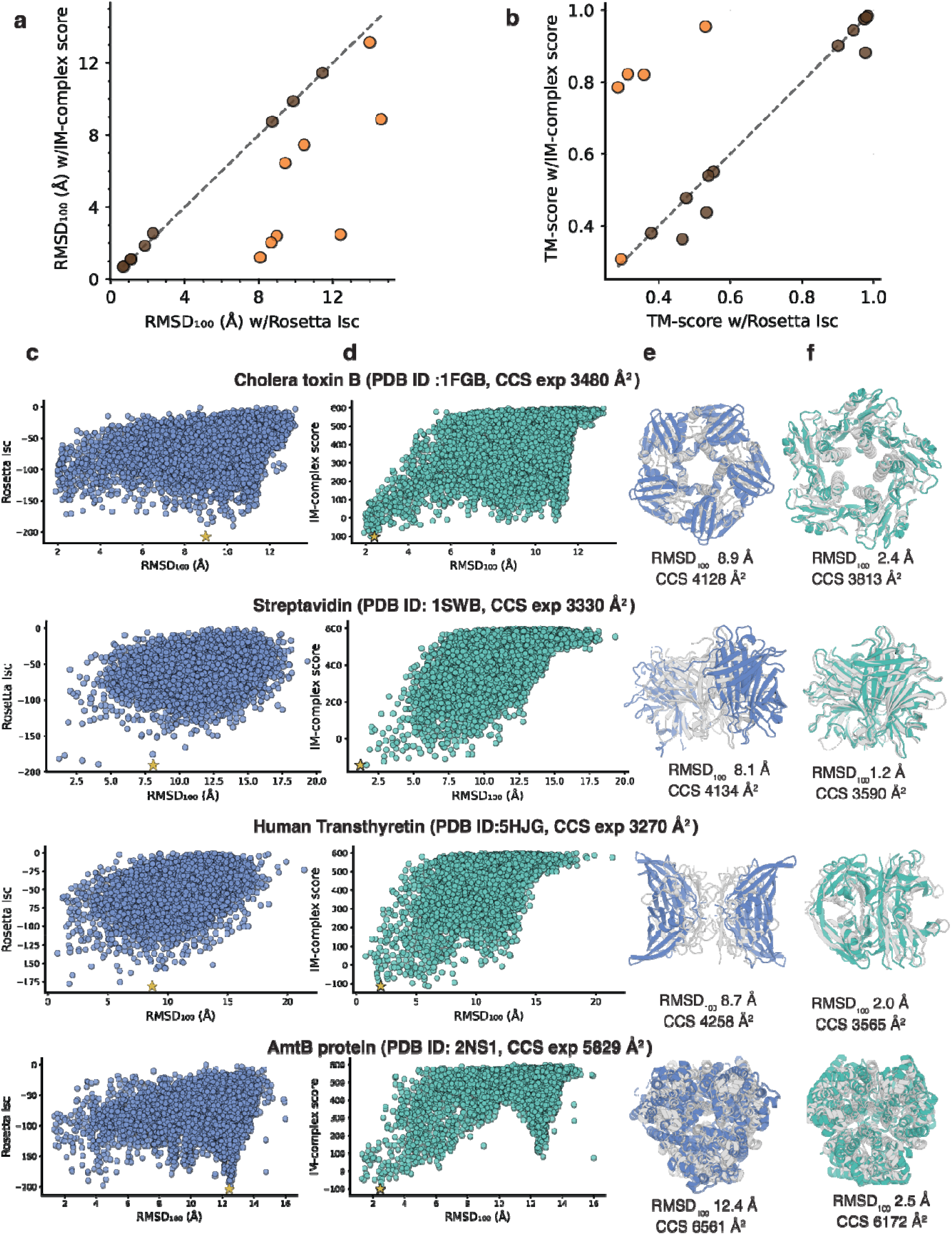
Improved model selection was observed when using the IM-complex score for the 17 proteins in the benchmark dataset. The predicted models from the IM-complex score were compared to those scored with the Rosetta Isc function in terms of their a) RMSD□□□ (root mean square deviation normalized by sequence length) and b) TM-Score (template modeling score). The protein complex structures that improved upon using the IM-complex score are colored orange, while the remaining proteins are colored brown. c-f) Comparison of modeled structures of four significantly improved proteins with their experimental CCS values. Respective protein score vs RMSD□□□ plots scored with c) Rosetta Isc (blue scatter plot) and d) IM-complex score (green scatter plot). The best scoring model is marked with a yellow star in each plot. Corresponding best scoring structures generated with e) the Rosetta Isc function (blue) and f) the IM-complex score function (green) are compared to their native structures (gray) and respective RMSD□□□ and PARCS-complex CCS values are listed.

Four protein complexes (cholera toxin B (PDB ID: 1FGB, homo 5-mer), streptavidin (PDB ID: 1SWB, homo 4-mer), human transthyretin (PDB ID: 5HJG, homo 4-mer), and AmtB protein (PDB ID: 2NS1, homo 3-mer)) showed significant improvement, with RMSD□□□ values decreasing from greater than 8 Å to less than 3 Å. This improvement is shown in Figure 5c-f. Although Rosetta generated near-native poses, the Rosetta Isc often failed to rank them correctly (Figure 5c). In contrast, the shape and size information captured by experimental IM-MS data guided the IM-complex score toward more native-like structures (Figure 5d). Figures 5e and 5f further highlight the improvement, showing how the best-scoring structures shifted from those selected by the interface score to those best scored by the IM-complex score. These results demonstrate that experimental IM-MS CCS measurements provide valuable complementary information, enhancing the ability to identify correct quaternary arrangements from docking ensembles.

## Conclusion

Ion mobility mass spectrometry can preserve near-native protein structures in the gas phase, is fast, requires minimal sample, and provides valuable structural insights into the global shape and size of protein complexes. However, this information alone is sparse and insufficient for full structure determination. Deep learning networks such as AF have been successful in predicting monomer and complex structures, but still struggle with modeling many proteins, especially multi-subunit complexes. We developed a method that leverages the complementary strengths from deep learning and physics-based modeling and addresses the limitations of current state-of-the art protein complex structure prediction.

For protein complexes, the original PARCS method underestimated CCS values, so we reparametrized it using TJM CCS values, reducing percent error from 8.1% to 3.9%. We then incorporated CCS into Rosetta docking through an IM-complex score that penalizes deviations between predicted and reference CCS. For the ideal dataset (challenging cases where AFM often misassigned symmetry, connectivity, or interfacial contacts despite accurately predicted monomers), the IM-complex score proved beneficial in improving structure quality compared to just modeling with AFM or Rosetta Isc. Docking AF subunits with Rosetta and rescoring with the IM-complex score produced models with more correct global shape, symmetry, and interface geometry. Overall, compared to AFM, the IM-complex pipeline improved 68.4% of complexes with an average reduction in RMSD of 10.2 Å for the improved protein complexes. Using experimental CCS values to rescore showed similar trends as well. In a benchmark set of 17 complexes, when experimental CCS was used to rescore, the average decrease in RMSD was 11.2 Å for the improved protein complexes (8/17). These results highlight the broader value of integrating CCS measurements into structure prediction and docking workflows. The detailed description of methods, as well as a tutorial is provided in the Methods and Supplementary Information.

In this study we tested whether the shape and size information encoded in a single CCS value could help distinguish near-native models from incorrect ones for protein complexes. Our results showed that including CCS values improved structure prediction in several instances. However, there are cases that showed no or little improvement and generating 10,000 models remains computationally intensive in Rosetta. To address this issue in the future, we plan to incorporate CCS values directly into the AF network to leverage even larger improvement in structure prediction. We also aim to extend this work by incorporating additional experimental restraints, such as cryo-EM, SID and NMR data, to develop a more comprehensive integrative modeling framework.

The developed PARCS-complex application and IM-complex scoring application is freely available within Rosetta (GitHub link: https://github.com/RosettaCommons/rosetta/).

## Methods

### Datasets

#### Reparameterization Dataset

To renormalize the original PARCS algorithm using TJM CCS values, a dataset of 461 protein complexes was curated from the RCSB PDB. The selected structures had refinement resolutions between 0.5 and 1.5 Å and molecular weights ranging from 5 kDa to 300 kDa (see Figure S3), encompassing a broad spectrum of complex sizes. The PDB IDs of the 461 proteins are reported in Dataset S1. The dataset consisted of 390 homomers and 71 heteromer including 298 dimers, 28 trimers, 96 tetramers, 14 pentamers, 16 hexamers, 1 heptamers, 2 octamers, 3 decamers and 4 dodecamers.

#### Benchmark Dataset

To validate the IM-complex score for protein complexes with experimentally measured CCS values, a benchmark dataset was compiled with CCS values reported in the literature.^68–73^ For each protein complex, the CCS corresponding to the lowest charge state was used as representative experimental CCS (see Table S3). The dataset consisted of 14 homomers and 3 heteromers including 5 dimers, 1 trimer, 7 tetramers, 2 pentamers and 2 hexamers.

#### Ideal Dataset

The ideal dataset was designed to test how valuable CCS IM-MS data is in improving protein structure prediction, assuming there was no error in predicting CCS values. We assembled a challenging dataset where AFM performed poorly. It consisted exclusively of protein complexes whose AFM RMSD to the native structure was greater than 4 Å. The dataset consisted of 36 homomers and 3 heteromers including 23 dimers, 1 trimer, 8 tetramers, 4 pentamers and 2 hexamers. These proteins did not have experimental CCS values. Hence, the simulated PARCS-complex CCS value of the native structure was used as a stand-in for the experimental CCS.

### Reparameterization of PARCS

To improve the accuracy of PARCS and enable more reliable CCS predictions for protein complexes, we established a relationship between TJM Collidoscope CCS values and PARCS CCS values. To assess this relationship, we computed CCS values for each protein complex in the reparameterization dataset using the PARCS algorithm (PARCS) and obtained TJM CCS values with Collidoscope.

#### CCS calculation with the TJM method

We calculated CCS values using Collidoscope for each of the 461 protein complexes in the reparameterization dataset. To generate representative structures for TJM calculations, vacuum molecular dynamics (MD) simulations were performed using GROMACS^74^ with the GROMOS 43a2 force field. Following the Collidoscope workflow,^64,65^ each protein complex was first energy-minimized in a water environment, then subjected to a brief (1 ns) MD relaxation in water to ensure proper folding. After this step, water molecules were removed, and the proteins underwent a 10 ns equilibration run and a 10 ns production run in the NVT ensemble at 300 K. CCSs for representative structures were computed using He buffer gas in the Collidoscope Trajectory Method.^64^

#### CCS calculation with the PARCS method

CCS values were calculated for the PDB structures of each protein in the reparameterization dataset using the PARCS application in Rosetta,^50^ employing 330 random rotations and a probe radius of 1.0 Å to simulate helium gas conditions.

#### Reparameterization

The reparameterization Eqn. 1 was obtained by fitting a power-law model of the form y =kx^a^ to all 461 protein complex CCS values using nonlinear least-squares regression. The parameters a and k were optimized by minimizing the sum of squared differences between Collidoscope TJM CCS and PARCS CCS using the curve_fit function in SciPy.^75^

### IM-MS complex score

PARCS-complex CCS values provide valuable information about the overall global size, shape, and inter-subunit arrangement of the complex. To make use of this information, we developed an IM-complex score term that quantifies how well a predicted protein complex agrees with experimental IM-MS data. The IM-complex score term is combined with the standard Rosetta Isc (from the REF2015 energy function) to generate a final IM-complex Score (see Eqn. 2).

The method uses a penalty function (see Eqn. 3, 4) (see Figure S4) that penalizes models based on their absolute CCS deviation (ΔCCS) from the experimental value. Models with larger deviations receive higher penalties, while those closer to the experimental CCS receive lower/no penalties. Compared to a similar function we used for monomeric proteins^50^ (with bounds of 0 and 100) this function employs higher lower bound (LB) and upper bound (UB) values. The LB of 60 and UB of 1350 were chosen to account for the larger size and greater variation in CCS values of protein complexes creating a custom penalty function specifically for these multimeric assemblies. This score is implemented in Rosetta and acts as a restraint during structure selection, helping to guide the identification of native-like assemblies.

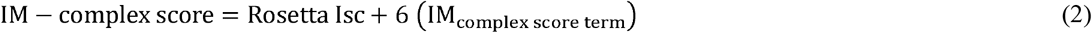

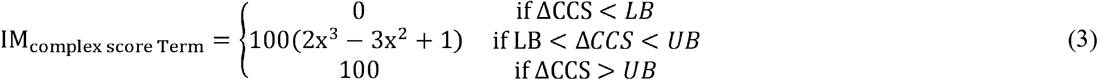

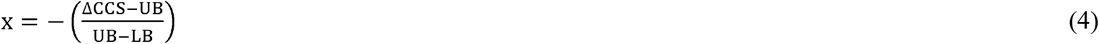

### Docking Protocol

#### Subunit generation

Subunits were generated using AlphaFold v2.2. The --max_template_date flag was set to 1900-01-01 to ensure a consistent template cutoff across all proteins and to confirm that our method remains applicable even when no templates are available. The --model_preset option was set to monomer for single chain AF or multimer for multi chain AFM predictions.

#### Ensemble generation

To better account for conformational variability at the interfaces while retaining the structures close to those predicted by AF or AFM, we generated subunit ensembles by perturbing the backbone structures using four Rosetta backbone movers: backrub,^76^ relax,^77^ normal mode analysis (NMA),^78^ and shear.^79^ Ten structures were generated with each mover, resulting in a pool of 40 perturbed backbone conformations per subunit.

#### Docking method

The ensembles generated from subunits were docked with the Rosetta Docking protocol^40,80^. We generated 10,000 models per protein complex to ensure broad conformational sampling. The docking method is described in the Supplementary tutorial. The models were then evaluated using both the Rosetta Isc and our newly developed IM-complex score, which integrates shape and size information derived from IM-MS measurements. For the ideal dataset, the PARCS-complex CCS prediction of the native structures was used as the reference CCS value. Physically unrealistic structures among the docked models were removed by identifying models above a minimum interchain distance of 25 Å between any Cα atoms within the respective chains. This was implemented using a breadth-first search (BFS) algorithm, where nodes represented chains and edges represented minimum interchain distances.

### AFM calculations

Protein complex structure was predicted for comparison using AlphaFold v2.2. The --max_template_date flag was set to 1900-01-01 to ensure a consistent template cutoff across all proteins and to confirm that our method remains applicable even when no templates are available. The --model_preset option was set or multimer for multi chain AFM predictions and the flag –reduced_database was used. pLDDT values were calculated from b factor column of the predicted structures. pTM and ipTM values were obtained from pickle files associated with each predicted structure.

### Analysis Metrics

We evaluated the modeled protein complexes generated by AFM and Rosetta using multiple structure quality metrics. The root-mean-square deviation (RMSD) was computed in PyMOL, while the RMSD□□□^81^ (Normalized RMSD, see definition in SI) was derived from RMSD and sequence length to account for protein size differences across the datasets. Additionally, the template modeling score (TM-score) was calculated using USalign^82^ to assess overall structural similarity, and the DockQ^83^ score was used to evaluate the quality of the docking interface.

A tutorial with all the commands used during the IM-complex docking protocol is provided in the Supplementary information. More information about the MD simulations and BFS algorithm can be found on the LindertLab GitHub repository (https://github.com/LindertLab/IM-complex).

## Supporting information

Figure S1,2,3,4, Table S1,2,3, Dataset S1, Tutorial

## Acknowledgements

We thank fellow Lindert lab members for their valuable discussions and guidance. We extend our gratitude to the Rosetta Commons community for thoughtful discussions during RosettaCon. We also acknowledge the computational resources provided by the Ohio Supercomputer Center (OSC)^84^. Additionally, this work used computational and storage services associated with the Hoffman2 Cluster which is operated by the UCLA Office of Advanced Research Computing’s Research Technology Group. This work was supported by RM1-GM149374.

## Notes

### Competing Interest Statement

The authors have declared no competing interest.

