## Supplementary material for "Ion Mobility Mass Spectrometry Guided Modeling with AlphaFold and Rosetta Improves Protein Complex Structure Prediction": Figure S1,2,3,4, Table S1,2,3, Dataset S1, Tutorial

Figure S2………………………………………………………………………………………….3

Table S1…………………………………………………………………………………………...4

Table S2…………………………………………………………………………………………...6

Table S3…………………………………………………………………………………………...7

Figure S3.……………………………………………………………………………………….….8

Figure S4.……………………………………………………………...…………………………...9

Dataset S1……………………………………………………………….................................…...10

Tutorial……………………………………………………………………………………….........11

**Figure S1**: Distribution of normalized radius of gyration change (R_g_normalized = (R_g_model − R_g_native ) / R_g_native) of 461 protein complexes in the reparameterization dataset.

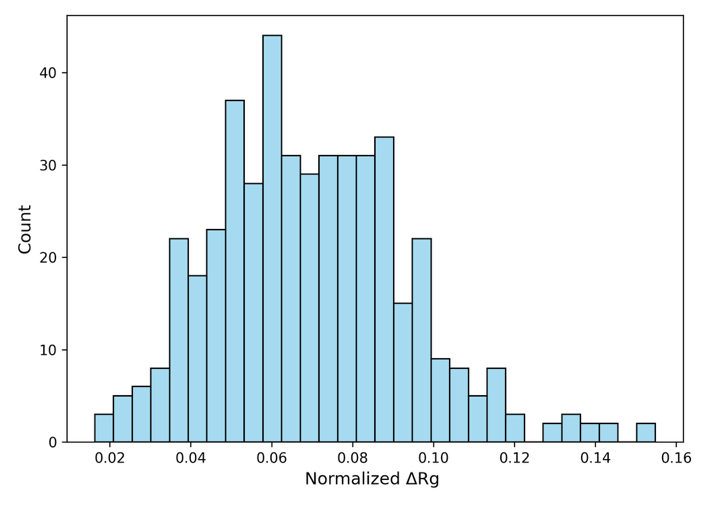

**Figure S2. Quadrant analysis of AFM confidence metrics and structural accuracy for ideal dataset proteins.**
(a) mean pLDDT versus RMSD of AFM-predicted structures relative to native PDB conformations.
(b) Predicted model confidence score (0.8 × ipTM + 0.2 × pTM) versus RMSD relative to native PDB structures.

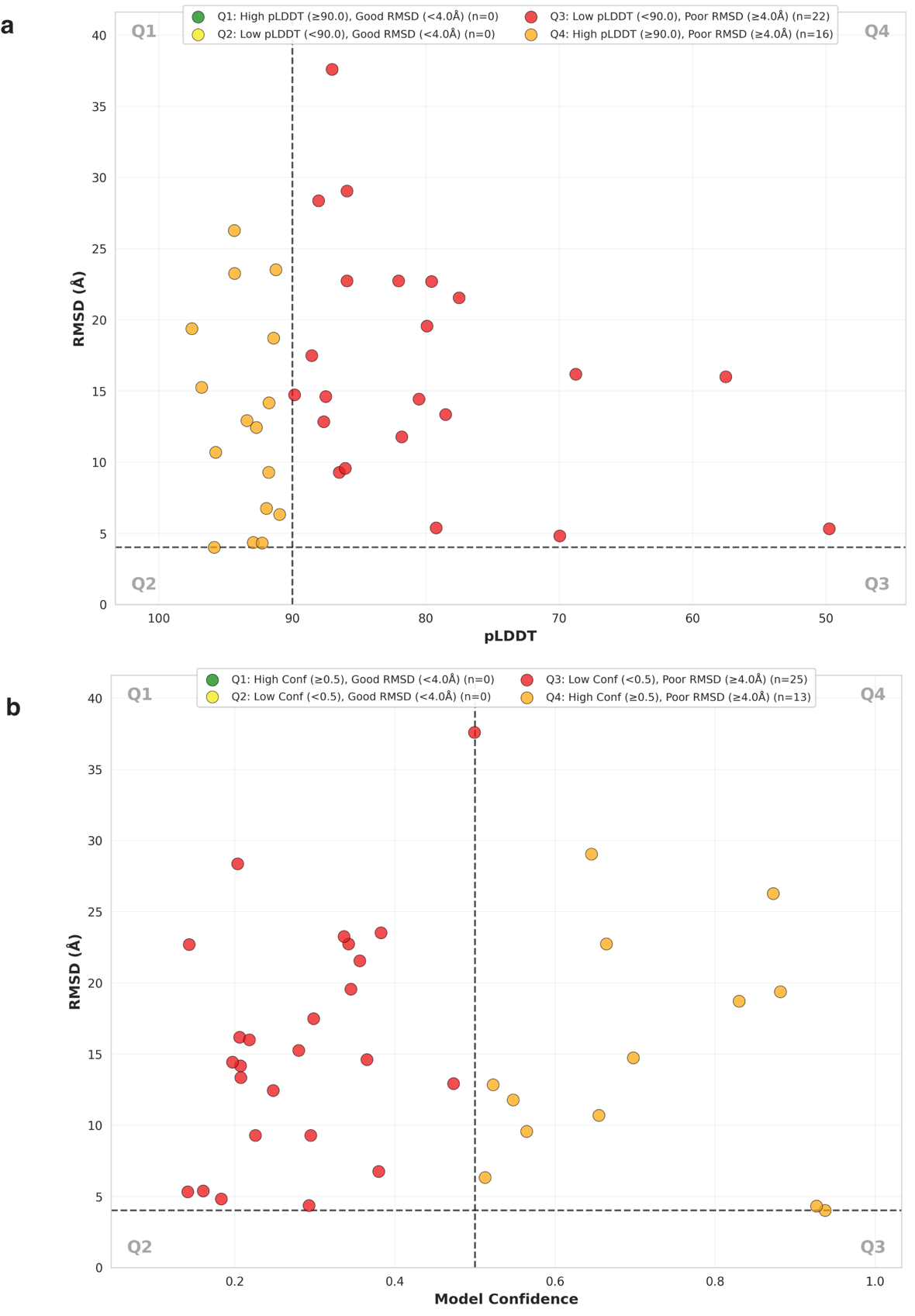

**Table S1:** Summary of protein complexes in the ideal dataset showing their PDB ID, protein name and average RMSD of each monomeric chains of the predicted subunit relative to the experimental structure.

| **PDB ID** | **Protein name** | **oligomeric state** | **Average RMSD of each unique monomeric chain of the predicted subunit(s) [Å]** |
| --- | --- | --- | --- |
| 1FXY | coagulation Factor Xa-Trypsin | 2 | 2.1 |
| 2CH6 | human N-acetylglucosamine kinase | 2 | 2.4 |
| 2DV9 | peanut lectin | 4 | 0.65 |
| 2IPJ | h3alpha-hydroxysteroid dehydrogenase type 3 mutant Y24A | 2 | 0.6 |
| 3BX7 | Engineered human lipocalin 2 | 4 | 0.8,1.25 |
| 3H45 | Glycerol kinase | 2 | 1.3 |
| 3T6O | Anti sigma factor antagonist (STAS) domain protein | 6 | 0.79 |
| 4D3Q | Point mutated DUSP19 | 2 | 0.5 |
| 4DYM | ACVR1 kinase domain | 2 | 1.7 |
| 4FN9 | Ancestral 3-keto steroid receptor | 2 | 0.7 |
| 4JCF | S268F Variant of JC Polyomavirus Major Capsid Protein | 5 | 0.6 |
| 4NPU | HIV-1 Protease Multiple Mutant P51 | 2 | 1.6 |
| 4WLE | Citrate bound MDH2 | 4 | 1.0 |
| 5D65 | Macrophage inflammatory protein | 5 | 1.3 |
| 5HGG | Camelid derived antibody fragment | 4 | 0.67, 2.45 |
| 5INX | A C69-family cysteine dipeptidase | 5 | 0.7 |
| 5XTJ | Mannanase | 2 | 0.5 |
| 6R5Z | 9-bladed beta-propeller | 3 | 1.3 |
| 6TAR | dArc1 capsid | 5 | 2.0 |
| 1JLT | Vipoxin complex | 2 | 0.9, 1.4 |
| 4JEA | engineered Zn-RIDC1 construct with four interfacial disulfide bond | 4 | 0.6 |
| 4YW6 | Galactophilic lectin | 4 | 0.8 |
| 7MK4 | engineered cyt cb562 variant, DiCyt2 | 2 | 1.2 |
| 7MNK | tetramerization element of NUP358/RanBP2 | 4 | 0.5 |
| 7N6V | HIV-1 protease | 2 | 1.3 |
| 7QGW | Human Cathepsin L2 | 2 | 0.8 |
| 7TC5 | All Phe-Azurin variant - F15Y | 2 | 0.4 |
| 7TO5 | HIV-1 wild type protease | 2 | 1.1 |
| 7W70 | HIV-1 wild type protease | 2 | 0.8 |
| 8AVU | Aureocin A53 complex | 2 | 0.4 |
| 8DCH | HIV-1 protease Clinical isolate | 2 | 1.4 |
| 8F0F | HIV-1 wild type protease with GRL-110-19A | 2 | 1.2 |
| 8FUI | HIV-1 wild type protease with GRL-02519A | 2 | 1.2 |
| 8FUJ | HIV-1 wild type protease with GRL-03419A | 2 | 1.2 |
| 8VIJ | Truncated hemoglobin | 4 | 0.6 |
| 8X6P | Isomerase protein | 2 | 0.4 |
| 8Z28 | FK506-binding protein | 2 | 0.5 |
| 6E2B | Ubiquitin complex | 6 | 1.1 |

**Table S2**: Summary of protein complexes in the benchmark dataset showing their PDB ID, protein name and average RMSD of each monomeric chains of the predicted subunit relative to the experimental structure.

| **PDB ID** | **Protein name** | **oligomeric state** | **Average RMSD of each unique monomeric chain of the predicted subunit(s) [Å]** |
| --- | --- | --- | --- |
| 1AQF | Pyruvate kinase | 4 | 1.2 |
| 1AVE | Avidin | 4 | 1.0 |
| 1EM8 | DNA POL III | 2 | 1.2, 1.1 |
| 1FGB | Cholera Toxin B | 5 | 1.8 |
| 1GNH | Human C reactive protein | 5 | 0.5 |
| 1HK9 | RNA binding Hfq protein | 6 | 0.4 |
| 1SWB | Streptavidin | 4 | 0.4 |
| 1WBJ | Tryptophan synthase | 4 | 0.9 |
| 2AIY_AB | Dimer form of human insulin | 2 | 1.3, 5.1 |
| 2INC | toluene/o-xylene Monooxygenase Hydroxylase | 6 | 0.67, 0.52, 0.51 |
| 5ENV | Carboxyltransferase Subunit of ACC from Escherichia coli | 4 | 0.7 |
| 5HJG | Human Transthyretin | 4 | 0.7 |
| 6QI6 | recombinant bovine beta-lactoglobulin | 2 | 1.7 |
| FRAB | Salmonella deglycase | 2 | 2.7 |
| 1VLN | Concanavalin A | 4 | 1.4 |
| 1BSY | Bovine Beta-lactoglobulin | 2 | 2.0 |
| 2NS1 | Amtb protein | 3 | 0.5 |

**Table S3**: Experimental CCS values (in He buffer gas) of the protein complexes in the benchmark dataset.^1-6^

| **PDB ID** | **Experimental CCS value [Å^2^]** |
| --- | --- |
| 1AQF | 9920 |
| 1AVE | 3500 |
| 1EM8 | 2520 |
| 1FGB | 3480 |
| 1GNH | 6400 |
| 1HK9 | 3341 |
| 1SWB | 3330 |
| 1WBJ | 7629 |
| 2AIY_AB | 757 |
| 2INC | 9963 |
| 5ENV | 6940 |
| 5HJG | 3270 |
| 6QI6 | 2850 |
| FRAB | 3985 |
| 1VLN | 5360 |
| 1BSY | 2670 |
| 2NS1 | 5829 |

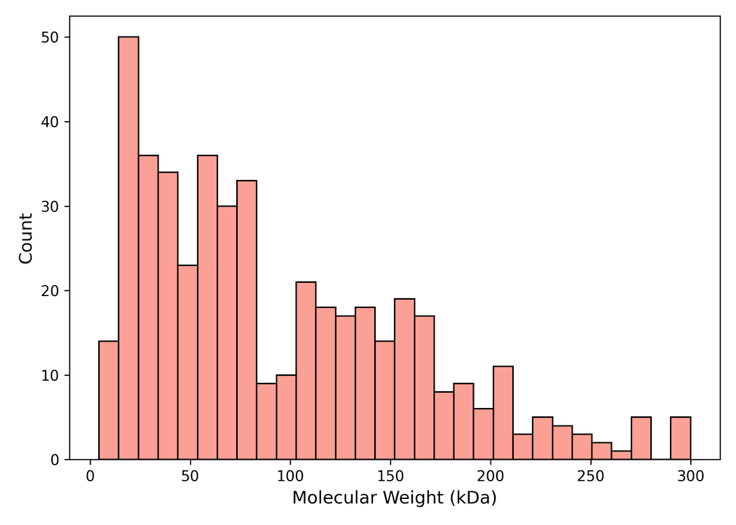
**Figure S3:** Molecular weight distribution of the 461 protein complexes in the reparameterization dataset.

**Figure S4: Illustration of the ion mobility score term (IM-complex score term).** The IM complex score term is a fade function in which LB and UB denote the lower and upper bound cutoffs, set to 60 Å² and 1350 Å², respectively. This term penalizes structures based on the absolute difference between the experimental collision cross section (CCS) and the predicted CCS of the structure (see eqn. 3, 4).

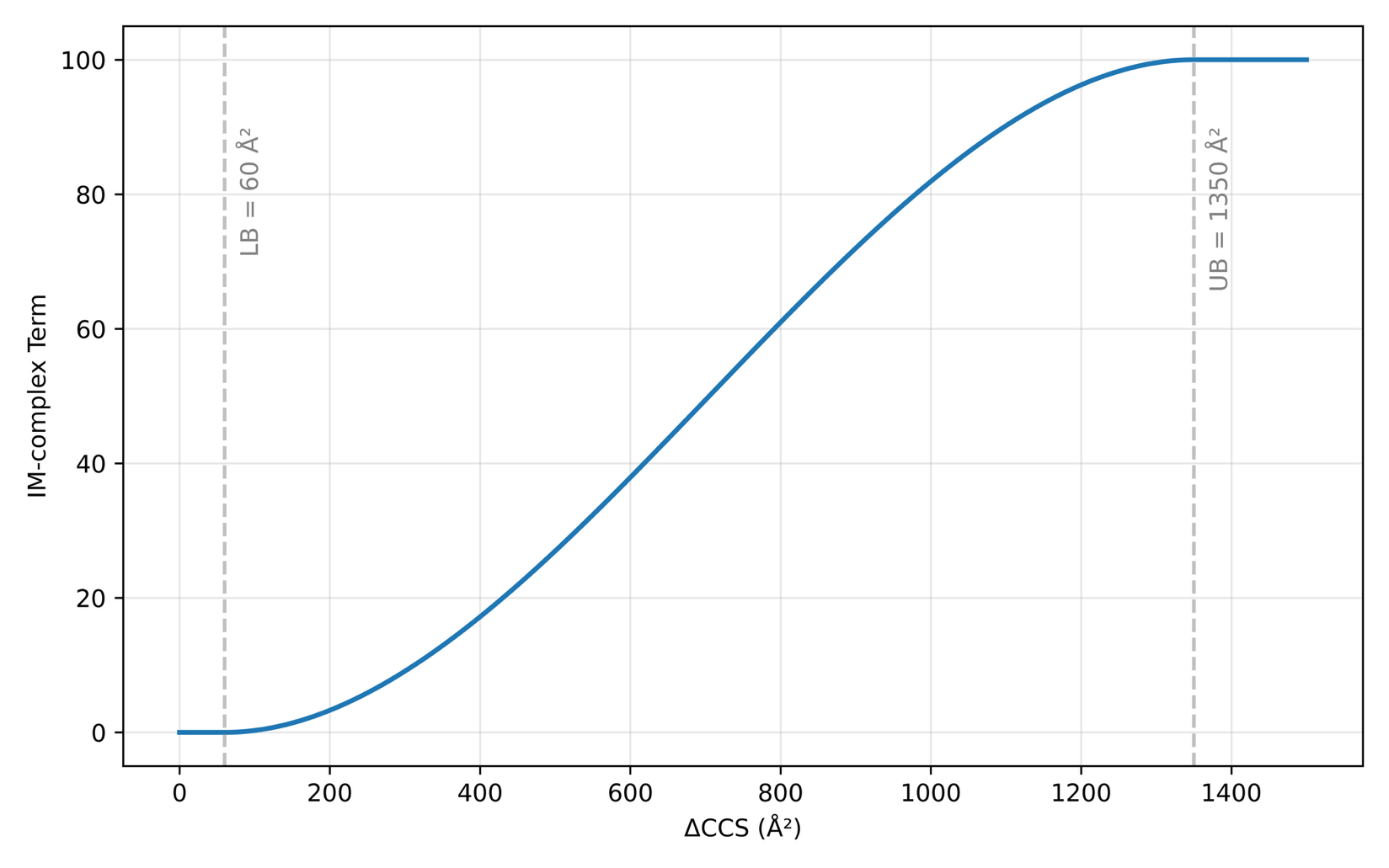

**Dataset S1:**

The PDB IDs of the protein complexes in the reparameterization dataset are listed below.

1B16, 1DJR, 1H2R, 1HET, 1JET, 1JR0, 1JUB, 1K3Y, 1KQP, 1LLR, 1MD2, 1MGO, 1N8K, 1N92, 1NEY, 1PZJ, 1QL0, 1RD9, 1RF2, 1SBY, 1U3W, 1WQW, 1WUH, 1X54, 2CVD, 2DN2, 2EGV, 2H8F, 2I7S, 2JHF, 2NXV, 2NYB, 2OB3, 2OLB, 2P4K, 2Q0L, 2R80, 2V8T, 2V9M, 2W72, 3A5F, 3A6R, 3BHL, 3BOM, 3D9A, 3E8M, 3F6B, 3L84, 3LJK, 3MJO, 3OQ6, 3TEM, 3W1A, 3W23, 3W75, 3W87, 3WA2, 4B2B, 4CW4, 4DWV, 4DXH, 4FP5, 4G41, 4HWW, 4HXQ, 4JEA, 4KEM, 4NFH, 4NG5, 4U7H, 4U9H, 4URE, 4UTU, 4XO6, 4Y59, 4YSL, 4YW6, 4ZUM, 5B5F, 5CDG, 5DGD, 5F5N, 5HEE, 5HH9, 5J3O, 5KCP, 5KJ1, 5LVS, 5LZG, 5MH2, 5VRK, 5YX1, 6CXX, 6E2B, 6E3D, 6E4D, 6EH4, 6FR3, 6H07, 6H5C, 6HCN, 6JQW, 6NNR, 6O91, 6OA7, 6OT4, 6OWM, 6SFE, 6SSF, 7E31, 7GRS, 7JY3, 7QGW, 7R66, 7UA6, 7UC9, 7UDD, 7UEC, 7UHV, 8BJ2, 8G39, 8G4V, 8HVL, 8JNW, 8OW6, 8VIJ, 8XAM, 8XDO, 9ART, 3A6Q, 2YEO, 7A2B, 5OLL, 6HS9, 6D0A, 6TXW, 5O3Y, 4JH2, 2YL0, 4DQE, 4EP4, 6Q00, 6DV0, 6MCR, 3ZTM, 7CAP, 4YIK, 7AEH, 5N20, 4JP2, 5MWX, 2YKZ, 6O48, 4J55, 4KTS, 3QP0, 6GMB, 6SDP, 5DWP, 5JVE, 2XI8, 5O6N, 6WNP, 3QIH, 6J9O, 4A7G, 7JQ2, 6IXD, 3NU3, 4ZIP, 4Z50, 6PRF, 6KAO, 7AWR, 7JOD, 4U8W, 5MAZ, 3QBF, 3S54, 4DFG, 4H30, 5DGU, 3K4V, 5B4T, 6H5G, 3PSM, 5I6K, 3ZQV, 6TAW, 3KFR, 3OXC, 5X2J, 6L5V, 5AH8, 6E9A, 5ESR, 3A2O, 6I42, 6DIF, 4CDA, 4GZG, 3VM9, 5JSL, 7NB9, 5BRY, 6U7O, 5YSX, 4NFL, 3PWM, 5JRA, 4E34, 6EP1, 3QAA, 6CDJ, 4CW2, 7KPH, 6WCO, 2Y5F, 6MV4, 3Q46, 5K1L, 4HE9, 4YHQ, 6E72, 7AQE, 3TOF, 3S43, 6YB7, 6T82, 4M7X, 3MMH, 4QBO, 4DWU, 6VVN, 7NIY, 6GSB, 3UVW, 6FJN, 6BLM, 6FVE, 6VCE, 7JR8, 5JFP, 4CX9, 7AOL, 6YQG, 5OF5, 3IJJ, 3LAS, 3M7U, 3ROF, 2WYT, 5JUA, 6OGL, 5JP7, 5JG1, 7K40, 3QGJ, 5AUR, 5LNI, 5LK9, 3PTL, 5JT4, 3I67, 5OK4, 4LUK, 6B38, 3L8W, 2WSB, 1W0D, 5ITW, 2YQU, 4WB7, 3T8I, 1OQS, 4H15, 1PBY, 3KF3, 1D4A, 4KEZ, 4O59, 1ZOS, 4EQR, 3X3X, 1FTR, 1GCO, 6FUR, 2INC, 1AVA, 1GD1, 2CFC, 6L8T, 3LQF, 1J31, 3KQR, 4K4B, 3DO6, 6FA1, 1FNU, 1WWS, 3KSE, 1M5S, 5VWO, 1R7A, 2VLF, 1DUV, 3LZ6, 3NTD, 1HG0, 6FQZ, 1XS1, 3X40, 1F5Z, 2CVP, 3RWB, 4WAS, 5A12, 1FD7, 1GEE, 4FN4, 1CCW, 1HRD, 2BBK, 4O63, 1EEF, 3WA3, 5GN2, 5IB0, 2QUL, 5ZP0, 4XR4, 3DAQ, 6H9E, 3SFW, 4DsG4, 3AI3, 4CZS, 6LCX, 6OIA, 6P73, 6Q9C, 6SPJ, 6T6P, 6V8C, 6VH9, 6YR3, 7B95, 7DY3, 7NNQ, 7PLJ, 7Q3X, 7QO8, 7QY9, 7QZ3, 7R65, 7WIR, 7YNH, 8BGX, 8G17, 8PYW, 1ADO, 1D7F, 1FBA, 1GQI, 1JQB, 1K3T, 1M3K, 1MG0, 1O9I, 1P1R, 1PU0, 1QDB, 1QH4, 1UFO, 1UM0, 1UXL, 1WXX, 1ZAH, 2BUH, 2C1D, 2D0I, 2O7C, 2ON5, 2OUI, 2R2N, 3BTO, 3EO8, 3FPC, 3HEA, 3IA2, 3IDS, 3LIN, 3NSX, 3RRS, 3RSY, 3T94, 3ZIW, 4O99, 4WUM, 4XLZ, 4ZFZ, 5D6O, 5F4X, 5GSM, 5LU0, 5MH3, 5VL0, 5VN1, 5VWQ, 5YHM, 5YMW, 6PTE, 6R8H, 6XT2, 7AGB, 7JQA, 7K35, 7UTC, 7WKL, 8AVL, 8GEK, 8PJ0, 1DM5, 1GK2, 1L6W, 1MTY, 1QH1, 2WYA, 3AI2, 3G5W, 3RGP, 4FO2, 4ZSW, 5CXK, 5DEI, 6F3M, 6G50, 6OP1, 7AHL, 7EFW, 7FJK, 7Q3T, 7TCS, 7TDL, 7V1Q, 7Y4G, 8OXF, 8XF2, 1H0H, 1KEK, 1KXQ, 2C3M, 2C42, 2WW2, 3S2E, 4ICM, 4IME, 4ISK, 4NG3, 4NI8, 8BH0.

**Tutorial**

This tutorial contains example Rosetta commands to run the calculations and scoring described in this manuscript.

**Usage of PARCS-complex to calculate CCS values for protein complexes**

To run PARCS-complex, a protein structure is given as input. CCS values are computed for one or more structures by calling the PARCS executable and pointing it to the Rosetta database. The executable is in the bin of Rosetta installation

<path/to/Rosetta>/main/source/bin/parcs_ccs_calc.default. <os><compiler>release \

-database <path/to/Rosetta>/main/database \

-in:file:s <structure> or -in:file:s <list of structures> \

-ccs_nrots <number_of_rotations> \

-ccs_prad <probe_radius_in_angstroms> \

-out:file:o <output_file_name>

**- ccs_multimer**

Flag and parameter description.

**-ccs_multimer** is the additional flag added to calculate CCS values for protein complexes.

path/to/Rosetta – root directory of the Rosetta installation

**os** – operating system tag (e.g. linux, mac)

**compiler** – C++ compiler used to build Rosetta (e.g. gcc, clang)

**structure** – input structure file for which CCS will be calculated

**number_of_rotations** – integer number of random orientations (default 300)

**probe_radius_in_angstroms** – buffer gas probe radius; 1.0 Å for helium

**output_file_name** – desired name of the CCS output file (if not given, CCS_default.out is used)

**Usage of the IM-complex score function to evaluate modeled structures**

This section describes how to evaluate Rosetta-generated models (from Rosetta docking pipelines) using IM-MS CCS data through the IM-complex score function.

Command to score a model looks like

<path/to/Rosetta>/main/source/bin/score.default. <os><compiler>release \

-database <path/to/Rosetta>/main/database \

**-in:file:s** <structure_from_prediction_protocol> or -in:file:l <for a list of models> \

**-ccs_nrots** <number_of_rotations> \

**-ccs_prad** <probe_radius_in_angstroms> \

-**ccs_exp** <experimental_ccs_data> \

**-score:**patch ccs_imms_complex.wts_patch

Flag and parameter description.

**score:patch** ccs_imms.wts_patch is the flag essential to score using IM-complex score function

path/to/Rosetta - Rosetta installation directory

**os** – Operating system identifier (e.g., linux, mac)

**compiler** – C++ compiler used to build Rosetta (e.g., gcc, clang)

**structure_from_prediction_protocol** – Model(s) produced by Rosetta ab initio or CM protocols

**number_of_rotations** – Number of random rotations used for CCS prediction (default: 300)

**probe_radius_in_angstroms** – Buffer gas probe radius (1.0 Å for He, reparametrization is done only in He gas environment)

**experimental_ccs_data** – IM-measured CCS value (Å²) for the protein under the same gas conditions

After successfully running the application, a score file will be generated. The column ‘ccs_imms_complex’ represents the weighted IM-complex score term. To obtain the IM-complex score, this term must be added to the interface score (column “I_sc”) from the score file, generated after docking with Rosetta.

**Example usage of creating decoys of cholera toxin protein (PDB ID: 1FGB, homo 5-mer)**

**with IM data**

To generate subunit for docking using AF.^7^

Obtain fasta sequence

wget https://www.rcsb.org/fasta/entry/1FGB

copy the fasta sequence of A chain to a separate file and name it 1FGB_A.fasta

Run AF

module load alphafold/2.2

run_alphafold.sh

--output_dir= <path/to/AF_subunit_output /1FGB> \

--fasta_paths=<path/to/1FGB_subunit_fasta_file> /1FGB_A.fasta \

--db_preset=full_dbs \

--model_preset=monomer \

--max_template_date=1900-01-01\

--use_gpu_relax \

To generate ensembles of subunits. (40 structures)

Relax

cd <path/to/docking_directory>/1FGB &&

<path/to/Rosetta>/main/source/bin/relax.linuxgccrelease \

-database <path/to/Rosetta> /database/ \

-in:file:s <path/to/AF_subunit_output /1FGB> /ranked_0.pdb \

-relax:quick \

-nstruct 10 \

-out:prefix af2_subunit_v22_relax_A_ \

Backrub

cd <path/to/docking_directory>/1FGB &&

<path/to/Rosetta>/main/source/bin/backrub.default.linuxgccrelease \

-database <path/to/Rosetta> /database/ \

-in:file:s <path/to/AF_subunit_output /1FGB>/ranked_0.pdb \

-nstruct 10 \

-backrub:mc_kt 0.6 \

-backrub:ntrials 200 \

-out:prefix af2_subunit_v22_backrub_A_ \

Shear

cd <path/to/docking_directory>/1FGB &&

<path/to/Rosetta>/main/source/bin/rosetta_scripts.linuxgccrelease \

-database <path/to/Rosetta> /database/ \

-in:file:s / <path/to/AF_subunit_output /1FGB>/ranked_0.pdb \

-nstruct 10 \

-parser:protocol shear.xml -out:prefix af2_subunit_v222_shear_A_ \

Shear.xml consists of:

<ROSETTASCRIPTS>

<SCOREFXNS>

<ScoreFunction name="r15_cart" weights="ref2015_cart.wts" />

</SCOREFXNS>

<RESIDUE_SELECTORS>

</RESIDUE_SELECTORS>

<TASKOPERATIONS>

</TASKOPERATIONS>

<FILTERS>

</FILTERS>

<MOVERS>

<Shear name="full_shear" scorefxn="r15_cart" temperature="0.2" nmoves="10" angle_max="360.0" preserve_detailed_balance="0"/>

</MOVERS>

<APPLY_TO_POSE>

</APPLY_TO_POSE>

<PROTOCOLS>

<Add mover="full_shear" />

</PROTOCOLS>

<OUTPUT scorefxn="r15_cart" />

</ROSETTASCRIPTS>

NMA

cd <path/to/docking_directory>/1FGB &&

<path/to/Rosetta>/main/source/bin/rosetta_scripts.linuxgccrelease \

-database <path/to/Rosetta> /database/ \

-in:file:s / <path/to/AF_subunit_output /1FGB>/ranked_0.pdb \

-nstruct 10 \

-parser:protocol nma.xml -out:prefix af2_subunit_v222_shear_A_ \

NMA.xml consists of:

<ROSETTASCRIPTS>

<SCOREFXNS>

<ScoreFunction name="r15_cart" weights="ref2015_cart.wts" />

</SCOREFXNS>

<RESIDUE_SELECTORS>

</RESIDUE_SELECTORS>

<TASKOPERATIONS>

</TASKOPERATIONS>

<FILTERS>

</FILTERS>

<MOVERS>

<NormalModeRelax name="nma" cartesian="true" centroid="false" scorefxn="r15_cart" nmodes="5" mix_modes="true" pertscale="1.0" randomselect="false" relaxmode="relax" nsample="1" cartesian_minimize="false"

/>

</MOVERS>

<APPLY_TO_POSE>

</APPLY_TO_POSE>

<PROTOCOLS>

<Add mover="nma" />

</PROTOCOLS>

<OUTPUT scorefxn="r15_cart" />

</ROSETTASCRIPTS>

All 40 perturbed structures then need to be listed by their full file paths in a single text file, with one structure per line, which is saved as ensemble_list_A. This file is then referenced as ensemble_list_A in the next step to collectively define and call the entire ensemble.

Docking the ensembles^8-13^

cd <path/to/docking_directory>/1FGB/ && <path/to/Rosetta>/main/source/bin/SymDock.linuxgccrelease @</path/to/docking_directory>/dock_flags

dock flags

-in:file:l <path/to/docking_directory>/1FGB/ensemble_list_A

-nstruct 10

-symmetry:perturb_rigid_body_dofs 5 60

-symmetry:symmetry_definition c5.symm

-symmetry:initialize_rigid_body_dofs

-symmetry:symmetric_rmsd

-packing:ex1

-packing:ex2aro

-out:prefix dock_

-out:path:all ./

-out:pdb

-out:file:fullatom

-out:file:scorefile score_file.sc

-detect_disulf false

-ignore_unrecognized_res

-evaluation:gdtmm

-evaluation:gdttm

-multiple_processes_writing_to_one_directory

AFM generation^14^. The following command was used to run AFM calculations

run_alphafold.sh \

--output_dir= <output_directory_path> \

--fasta_paths= <fasta_path> \

--db_preset=reduced_dbs \

--model_preset=multimer \

--max_template_date=1900-01-01 \

--use_gpu_relax \

--num_multimer_predictions_per_model=1

Dock Q calculation.^15^ DockQ <model>.pdb <native_structure>.pdb

TM-score calculation. TM score was calculated using USalign, ^16^

USalign <model>.pdb <native_structure>.pdb -mm 1 -ter 0 -TMscore 7 -cp

RMSD calculation. RMSD values for the C-alpha atoms were calculated in PyMOL with no outlier rejection (cycles=0). For homomeric complexes generated by AFM and Rosetta docking calculations, chain names could be interchanged in the output. To ensure the full symmetrical system is captured in the RMSD calculation, the chains were permutatively renamed, and the RMSD was calculated across all possible combinations with respect to the native PDB. The lowest RMSD value was then reported.

RMSD₁₀₀ calculation. RMSD values obtained from PyMOL were normalized with respect to the sequence length of the protein (N)(see Eqn. below).^17^

RMSD₁₀₀ = $\frac{RMSD}{1 + ln(sqrt(N/100))}$
